# Thyra: Bridging Mass Spectrometry Imaging and SpatialData for Unified Multi-Modal Analysis

**DOI:** 10.64898/2026.01.26.700792

**Authors:** Theodoros Visvikis, Wouter-Michiel Vierdag, Luca Marconato, Ron M.A. Heeren, Eva Cuypers

## Abstract

Mass Spectrometry Imaging (MSI) is a powerful technique for mapping molecular distributions, and its integration with other imaging modalities is crucial for comprehensive understanding of molecular systems. Fragmented data formats and the limitations of existing standards like imzML, challenge spatial biology centric multi-modal data analysis and adherence to FAIR data principles. This paper introduces Thyra, a modern Python library designed to convert MSI data into the SpatialData framework, a unified and extensible multi-platform file format that crucially integrates MSI into the broader spatial omics ecosystem. Thyra’s modular architecture, intelligent mass axis resampling, and sparse matrix backend address performance bottlenecks and facilitate seamless integration with tools for advanced spatial statistics and visualization. By adopting SpatialData, Thyra not only improves data interoperability, accessibility, and reusability, but also unlocks new avenues for multi-modal research, empowering scientists to integrate rich chemical information into diverse biological workflows.

## Main Text

Mass Spectrometry Imaging (MSI) allows for mapping molecular distributions across diverse samples, ranging from biological tissues to cultural heritage artifacts and advanced materials^1-3^. A unifying trend across these disciplines is the move towards multi-modal analysis, where MSI data is integrated with other imaging modalities to build a more comprehensive understanding of molecular landscapes^4^. This allows researchers to correlate the distribution of a drug in a tissue section with its underlying histology to assess spatial pharmacokinetics or map the degradation product of pigments in a historical painting to its high-resolution visible light image^5,6^. To create a truly comprehensive molecular picture, researchers increasingly need to integrate MSI with heterogeneous high-throughput technologies, such as histology, immunofluorescence, and spatial transcriptomics. However, the explosive growth in data volume and complexity has exposed a critical bottleneck: the lack of a standardized, community-driven format capable of robust multi-modal integration. The resulting fragmentation of data into proprietary vendor formats and the limitations of existing open standards impede the “FAIRness” (Findability, Accessibility, Interoperability, and Reusability) of MSI data, creating silos that restrict advanced integrative analysis^7^.

The current open standards, mzML^8,9^, and its imaging counterpart, imzML^10^, were significant innovations at their inception but predate the high-throughput demands of modern spatial biology^11^. Performance degradation in these formats is driven by the parsing overhead of XML-based metadata, which necessitates expensive tree-traversal operations that scale poorly with dataset size. Furthermore, these legacy formats lack support for critical modern features such as lazy loading, cloud-native access, and parallel read/write operations^12^. These limitations have driven many researchers to either custom formats^13^ – reducing interoperability – or towards vendor-specific solutions that rely on imzML for export, creating an inefficient conversion bottleneck. This process disincentivizes exploratory analysis, as the computational overhead of data preparation prevents novel hypotheses from being tested.

To avoid creating yet another isolated data silo for MSI, a sustainable approach is to align with the active communities of bioimaging and single-cell analysis by adopting the SpatialData framework^14^. Provided by the scverse ecosystem, SpatialData is a community-driven standard designed for high-resolution imaging and multimodal spatial molecular profiling. Built upon the OME-NGFF (Open Microscopy Environment Next-Generation File Format) open standard^15^, it uses Zarr and Apache Parquet as backend storage – two high performance, language-agnostic technologies used for saving tensors and tabular data, respectively. This provides a robust infrastructure with critical features for modern research: lazy loading, coordinate system transformations, spatial annotation, and cross-modal aggregation.

By connecting to this framework, MSI researchers can gain immediate access to the full suite of scverse analysis tools. Preliminary efforts, such as the *metaspace-converter* package^16^, have facilitated this connection by enabling the export of cloud-processed datasets from the METASPACE platform into SpatialData. However, this workflow is primarily designed to retrieve processed results – specifically molecular annotations and centroided spectra – rather than the full, continuous raw data required for de novo analysis or custom signal processing. Consequently, a gap remains for the direct, local conversion of complete raw vendor or imzML datasets independent of specific cloud pipelines. This lack of a standalone, general-purpose converter prevents researchers from leveraging the advanced analytical and multimodal integration capabilities that scverse offers^17^.

Here we introduce Thyra (from Greek θύρα, meaning “door” or “portal”) is a modern Python library for converting MSI data into the SpatialData format. By functioning as a robust and well-documented bridge, the library introduces MSI data into the collaborative world of modern spatial omics, enabling true cross-pollination between chemical and biological spatial analysis. This work provides a dual benefit: for the MSI community, it unlocks access to the mature scverse ecosystem, lowering the barrier for applying advanced spatial statistics and scalable visualization; for the broader spatial biology community, it makes the rich chemical information of MSI a readily accessible modality, allowing researchers to readily integrate spatial metabolomics or proteomics into existing multi-omics workflows (Fig 1).

**Figure 1.**
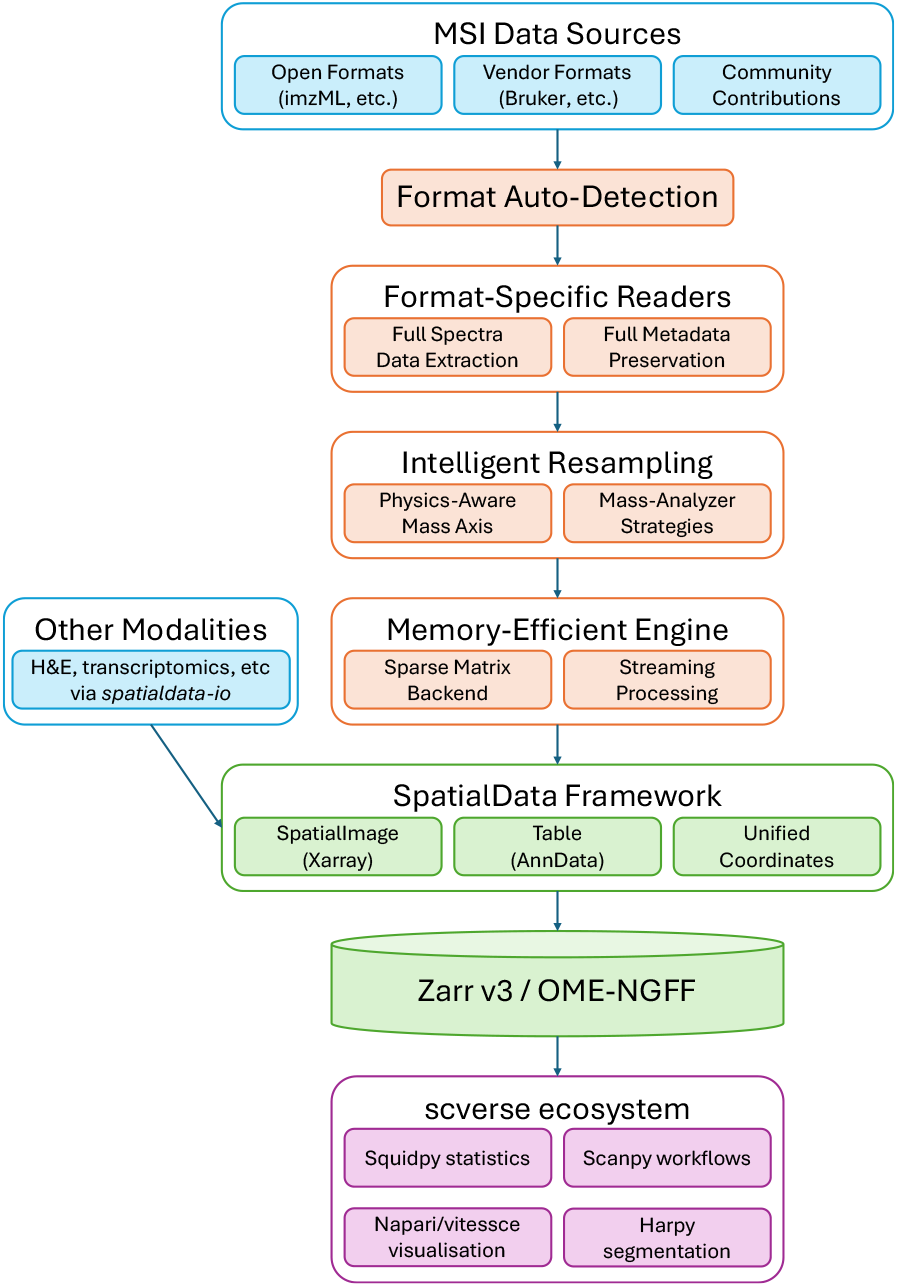
Overview of the end-to-end MSI data integration workflow. Diverse MSI inputs, including open (e.g., imzML) and proprietary vendor formats, are ingested via automated format detection to extract full spectra and metadata. The data undergo intelligent resampling using physics-aware strategies tailored to the mass analyser type, handled by a memory-efficient engine that utilizes sparse matrix backends and streaming processing. Additional spatial modalities (e.g., H&E, transcriptomics) can be integrated via spatialdata-io. The processed data are structured into the SpatialData model – incorporating SpatialImage (Zarr array), Table (Anndata), and unified coordinates – and stored using the cloud-optimized Zarr v3/OME-NGFF standard. This standardization enables downstream analysis and visualization within the scverse ecosystem using tools such as Squidpy, Scanpy, Napari, Vitessce, and Harpy.

## Results

### Improved storage efficiency and query performance

To evaluate the impact of the proposed method, we assessed improvements in storage efficiency. Standard approaches to unifying mass axes often lead to unmanageable file sizes; an attempt to convert our test dataset to a continuous imzML format resulted in a prohibitive 455 GB file. Our approach circumvents this via a sparse matrix backend. For the same dataset, the converted SpatialData object (4.83 GB) was substantially smaller than both the original Bruker .d file (21.2 GB) and the processed imzML exported from SCiLS (15 GB). This dramatic reduction is driven by the efficient, chunk-based compression native to the Zarr backend without compromising spectral fidelity, as confirmed by total ion current (TIC) comparisons (Fig 2).

**Figure 2.**
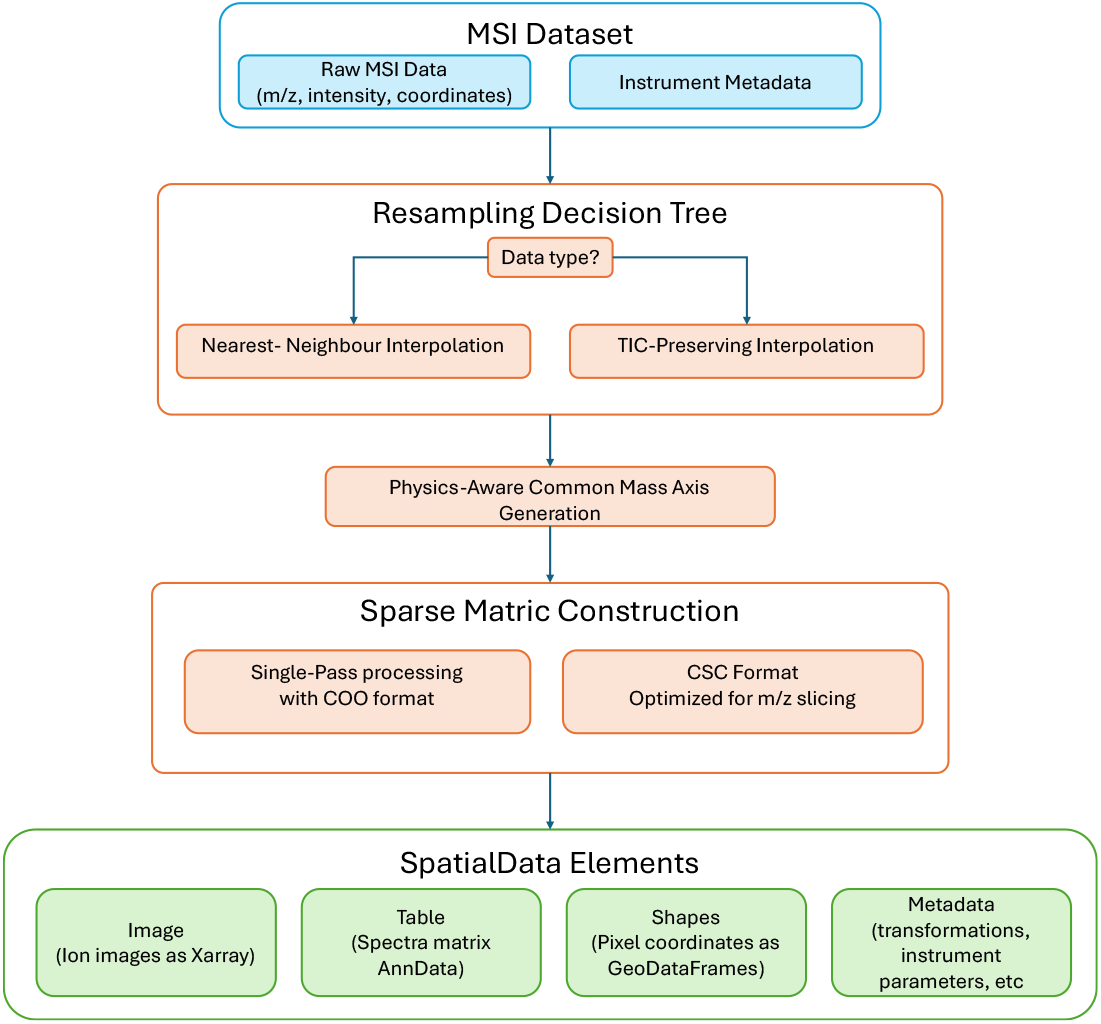
Detailed data processing and resampling workflow. Raw MSI data (m/z, intensity, coordinates) and instrument metadata are routed through a resampling decision tree that selects either nearest-neighbour or Total Ion Count (TIC)-preserving interpolation based on the acquisition type. This is followed by the generation of a physics-aware common mass axis to preserve spectral accuracy. A memory-efficient engine constructs the data matrix using single-pass Coordinate List (COO) processing, converting it to Compressed Sparse Column (CSC) format to optimize downstream m/z slicing. The final output is structured into distinct SpatialData elements: Image (ion images via Xarray), Table (spectral matrix via AnnData), Shapes (pixel coordinates via GeoDataFrames), and comprehensive metadata.

We further benchmarked performance on local storage, analysing initialization time and query latency (Fig. 3). The imzML format required 58 seconds to parse its large XML metadata – a step repeated for every session. In contrast, the acquisition-native Bruker .d format was fastest to initialise (0.5 s), while SpatialData presented a modest, one-time initialization cost of 8 seconds associated with loading the library ecosystem.

**Figure 3.**
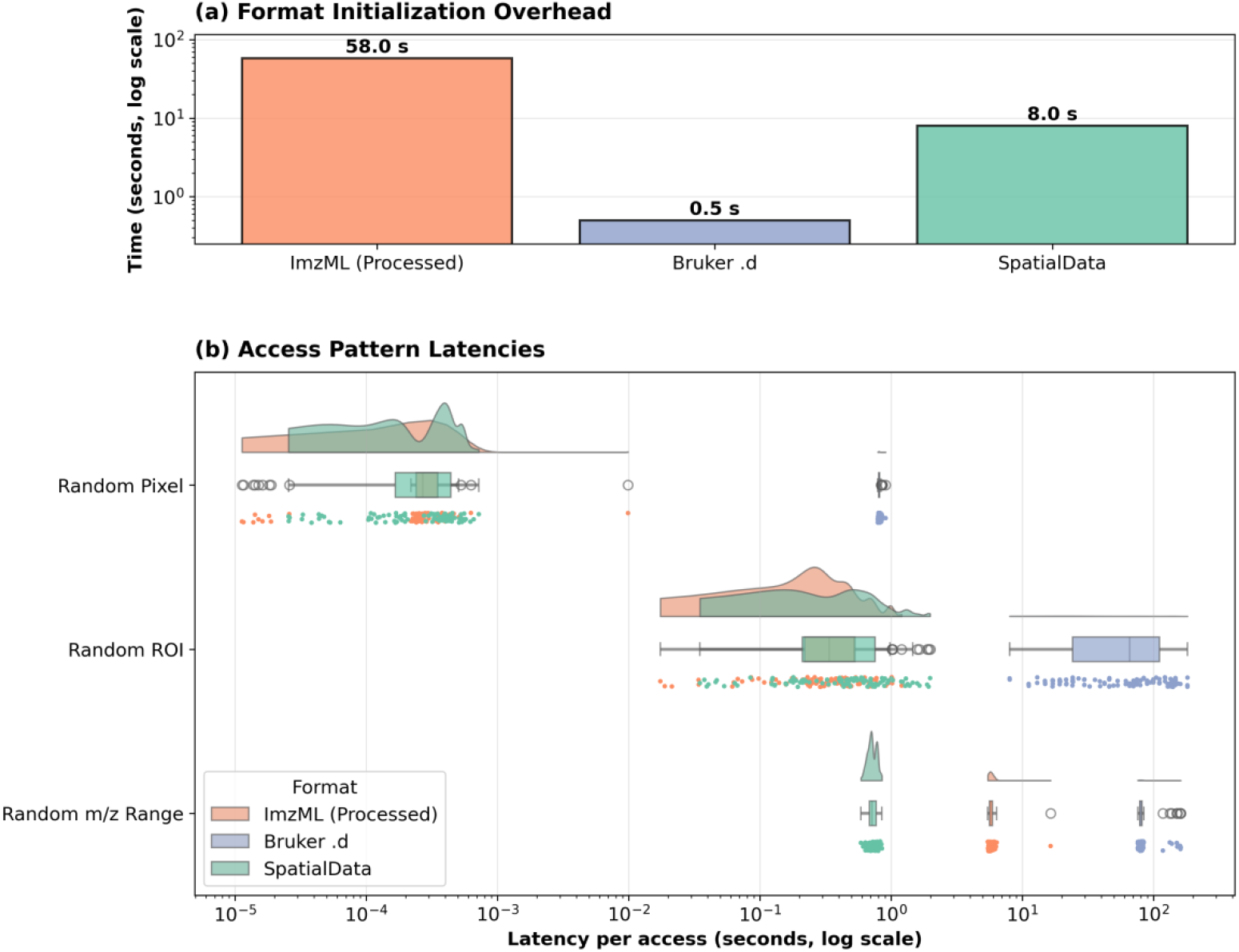
Performance Benchmark Results for Initialization Time and Query Latency. The benchmark compares three MSI data formats: the raw Bruker .d (blue), a processed imzML (orange), and the converted SpatialData object (green). (a) One-time initialization cost (“time to first query”). SpatialData (8.0 s) significantly outperforms imzML (58.0 s) in startup latency, while the raw Bruker format is fastest (0.5 s) (b) Raincloud plots displaying log-scaled query latency for 100 random queries across three distinct access patterns. For random pixel access, imzML retains a slight latency advantage, whereas for random ROI extraction, the performance between imzML and SpatialData is comparable. Crucially, SpatialData achieves an order-of-magnitude speedup for random m/z range queries—the bottleneck operation for generating ion images. The raw Bruker format shows consistently higher latency across all analytical patterns, confirming its unsuitability for direct downstream analysis.

In terms of query latency (Fig. 3b), the proprietary Bruker .d format was orders of magnitude slower for all analytical query patterns. For the critical task of querying by m/z range – the most common operation for generating ion images – the SpatialData format was nearly an order of magnitude faster than processed imzML. This performance gain arises because the Zarr backend allows for efficient slicing along the channel dimension, avoiding the costly non-contiguous read patterns required by row-major imzML files.

### Unified multi-modal spatial omics integration

The conversion to SpatialData immediately unlocks multi-modal capabilities (Fig. 4). We registered large, high-resolution MSI datasets to corresponding H&E histology images and Xenium transcriptomics data. While these distinct datasets initially exist in separate coordinate spaces, Thyra facilitates their alignment into a single, integrated SpatialData object.

**Figure 4.**
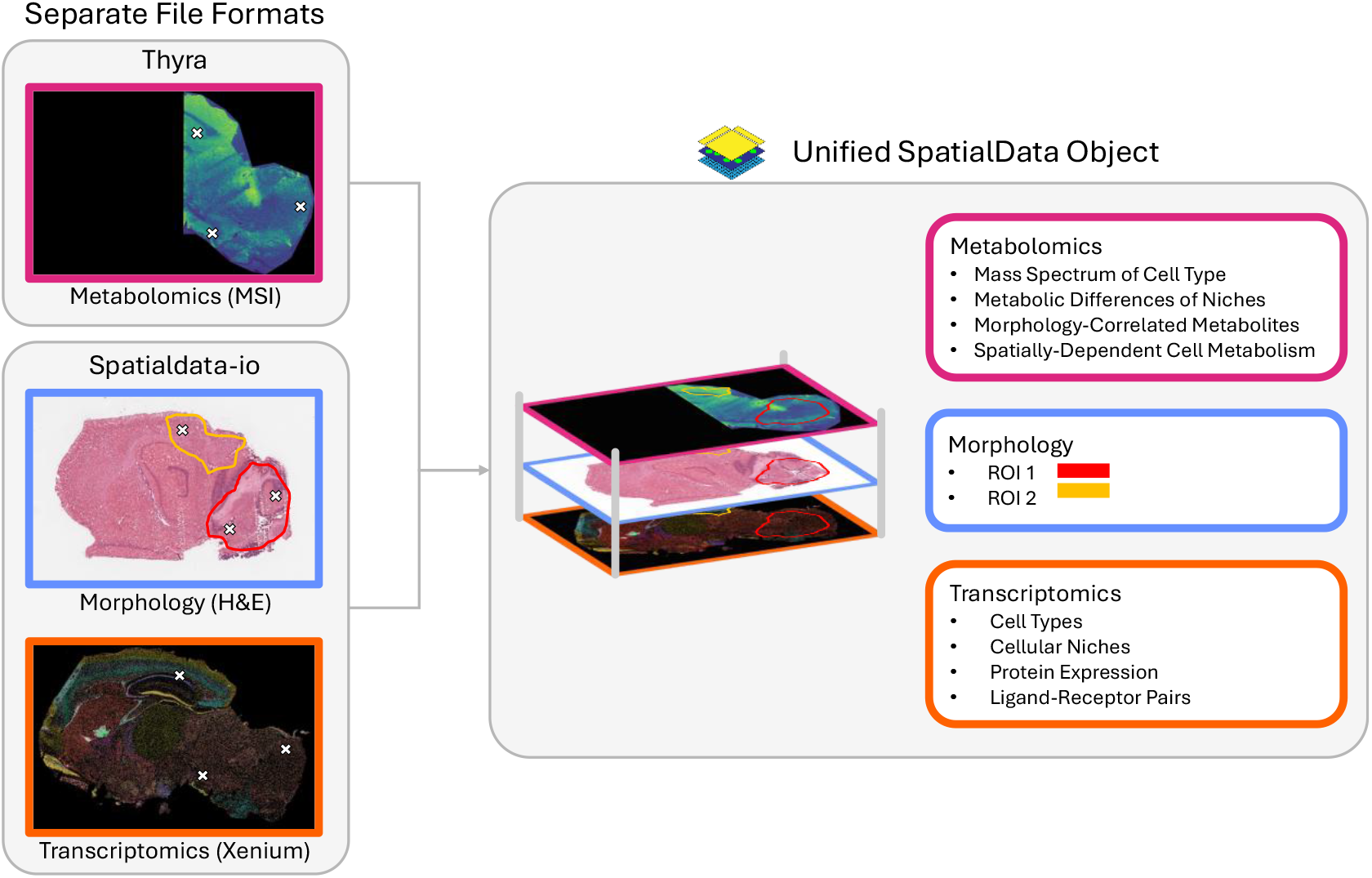
End-to-end workflow for aligning and leveraging diverse spatial omics modalities. Distinct input datasets, such as H&E histology (morphology), Xenium (transcriptomics), and MALDI-MSI (metabolomics), are registered into a common coordinate system using alignment tools such as EscDat^19^, resulting in a single, integrated SpatialData object. This unified container links aligned spatial layers with derived feature tables, enabling sophisticated downstream tasks: morphological feature extraction and cell segmentation (e.g., Segger)^20^, multi-modal analysis using squidpy, interactive, cloud-based visualization via Vitessce (facilitated by easy-vitessce) for quality control and dissemination. As a representative use case, biologically relevant Regions of Interest (ROIs) defined on H&E images can be used to spatially query aligned modalities, identifying metabolic profiles or cell type compositions that co-localize within specific histological regions. (Note: The H&E image displayed is a serial section and displayed as a representative example for illustrative purposes.)

This unified structure enables sophisticated downstream tasks. For example, biologically relevant ROIs can be defined directly from the H&E image’s morphological context to spatially query the aligned MSI modalities. This allows for the targeted extraction of molecular data, such as identifying dominant metabolic profiles or cell type compositions that co-localise within defined histological regions. Furthermore, the programmatic access provided by scverse libraries enables the application of spatial statistics algorithms (e.g., Squidpy) to correlate metabolic profiles with transcriptomic cell states^18^/

## Discussion

The rapid evolution of Mass Spectrometry Imaging into a cornerstone of multi-modal molecular imaging has created a critical need for data interoperability that existing standards were fundamentally not designed to meet. Our development of Thyra addresses this by providing a robust bridge between the established MSI landscape and the modern, collaborative ecosystem of SpatialData. This work goes beyond an engineering solution; it serves as a foundational step towards satisfying the scientific community’s commitment to FAIR data principles and unlocking the full potential of chemical imaging in spatial biology.

Historically, the fragmentation of MSI data into proprietary formats and the technical limitations of imzML have posed systemic barriers to FAIRness. Thyra improves this by adopting the OME-NGFF open standard, allowing MSI data to integrate seamlessly with other spatial omics modalities without customized parsers. The compact footprint of the resulting files directly improves accessibility and reusability, reducing storage costs and simplifying data transfer. Moreover, the cloud-native design of Zarr addresses a significant gap in the field: the need for large-scale and atlas-level centralised MSI databases. Because OME-NGFF/Zarr supports streaming data access (fetching only required chunks), it enables researchers to query and analyse massive datasets on remote resources without the need for full data transfer – a capability that has long powered the genomics revolution but remained absent in MSI.

This shift to a modern, chunked storage paradigm introduces specific performance trade-offs. Our benchmarks revealed that while SpatialData excels at analytical queries (like m/z ranges) essential for image generation, the processed imzML format retains a latency advantage for simple random pixel access on local storage. This is a known characteristic of the OME-NGFF/Zarr specification, where I/O overhead of accessing many small chunk files can impact local performance. Importantly, this limitation is already being addressed by the “sharding” feature in the Zarr v3 specification. Sharding groups multiple chunks into larger files, effectively mitigating the I/O bottleneck while retaining cloud-native benefits. Furthermore, our current benchmarks utilize default chunking strategies. Since Zarr performance is highly sensitive to the interplay between chunk size, compression, and file system block size, we anticipate that further optimization of these parameters—tailored to specific mass spectrometry data characteristics—will yield significant additional performance gains. We anticipate that future OME-NGFF standards incorporating sharding and appropriate parameters will rival or exceed traditional monolithic formats in local performance, effectively offering the best of both worlds.

Beyond technical metrics, the decision to adopt SpatialData provides the MSI community with immediate access to the mature scverse ecosystem, ensuring the technique does not become an isolated data silo. For the broader spatial biology community, our tool makes the rich phenotypic dimension of MSI a readily accessible modality. For instance, researchers can now leverage sophisticated featurization capabilities via cp_measure and squidpy to extract texture and morphology statistics^21^, perform segmentation using Harpy, and utilize interactive, cloud-native visualization tools like Vitessce^22,23^. For the broader spatial biology community, our tool makes the rich phenotypic dimension of MSI a readily accessible modality. Researchers can now add metabolomic and proteomic layers to transcriptomics studies to bridge the genotype-phenotype gap.

Finally, this standardization is a prerequisite for integrating molecular imaging into next-generation digital pathology workflows and clinical diagnostics. While the current implementation supports common open and vendor formats, its modular, plugin-based architecture is designed to catalyse community contributions. This community-centric design is pivotal for driving adoption; we are actively engaging with key partners, such as the scverse proteomics workgroup, to align development with the needs of related domains. This collaboration aims to establish a direct link to emerging spatial proteomics analysis tools, ensuring that the rich chemical data unlocked by MSI can be seamlessly integrated into dedicated downstream biological workflows. By fostering these connections, we lower the barrier for existing community efforts to adopt and build upon the software, further integrating MSI into the wider spatial omics landscape.

## Online Methods

### Software architecture and Design

Thyra is implemented as a modular, plugin-based Python library designed to decouple data ingestion, metadata interpretation, and format conversion. This architecture is managed by a thread-safe Registry System that performs automatic format detection and dispatches input files to the appropriate registered Reader plugin. To ensure extensibility, all plugins inherit from standardized abstract base classes (BaseMSIReader and BaseMSIConverter), allowing for the integration of new formats without modifying the core codebase. The software is distributed as an open-source project under the MIT license and is managed using Poetry for reproducible dependency resolution.

### Data processing & Conversion Workflow

- **Data Ingestion:** For imzML files, the parser ingests the dataset to extract spectral data and coordinates. For Bruker timsTOF data, the system interfaces directly with vendor SDKs, dynamically loading platform-specific shared libraries to access raw binary data while preserving instrument metadata.
- **Mass Axis Resampling:** Spectral harmonization is governed by a Resampling Decision Tree that parses instrument metadata to determine the acquisition type. Centroided data is processed using nearest-neighbour resampling to preserve peak fidelity, while profile data undergoes totalion-current (TIC) preserving interpolation. The system supports physics-aware non-linear axes to account for specific mass analyser characteristics (e.g., TOF vs Orbitrap).
- **Sparse Matrix Backend:** To handle datasets exceeding available RAM, the engine utilizes a Coordinate (COO) sparse matrix format. Arrays are pre-allocated based on metadata to prevent dynamic resizing. This avoids the computational overhead of repeated memory reallocation and data copying. Spectra are processed in a single pass: raw intensities are read, resampled, and non-zero values are inserted directly into the pre-allocated arrays. The final structure is converted to Compressed Sparse Column (CSC) format to optimize column-slicing performance during downstream analysis.

### SpatialData Mapping & Storage

The harmonized data is structured according to the SpatialData model. Ion images are stored as SpatialImage elements using Xarray, while spectral data and pixel-level metadata are organized into a Table using the AnnData standard. Pixel coordinates are translated into a unified physical coordinate system. The final object is serialized using the SpatialData format (an extension of the OME-NGFF specification evolving towards full compliance). This backend is supported by Zarr v3 storage, which supports sharding and asynchronous I/O for efficient cloud-native access.

### Validation & Implementation

Correctness is verified through a continuous integration framework comprising unit tests, and performance benchmarks. Conversion fidelity is validated via three quantitative metrics: metadata preservation (comparison of instrument parameters), data integrity (quantitative comparison of total ion current images), and spectral fidelity (direct comparison of individual mass spectra).

The software is managed using Poetry to ensure reproducible dependency resolution. It features a modular Command-Line Interface (CLI) built on the Click framework, incorporating automatic format detection, strict parameter validation, and verbose logging to facilitate pipeline integration. The package is hosted on the Python Package Index (PyPI) for standard deployment.

### Performance Benchmarking Strategy

To evaluate the performance characteristics of different mass spectrometry imaging data formats, we conducted a comprehensive benchmark using a single modest-scale dataset 18 GB raw, 918,855 pixels). We compared three formats: the original Bruker .d (vendor raw format), a processed imzML, and a SpatialData/Zarr representation. The two analysis-ready formats represent fundamentally different storage strategies: the processed imzML (exported from SCiLS) stores data in a ragged array mode with distinct m/z axes per pixel, while the SpatialData/Zarr version (generated via Thyra) stores all spectra against a single, common mass axis. All Benchmarks were executed sequentially on a single thread on a test environment consisting of an Intel Core i9-14900K (3.20 GHz), 128 GB RAM, and local SSD storage.

### Access Pattern Protocols

We designed three access patterns to stress-test data organization:

- **Random Pixel Access:** We generated 100 random pixel indices. For imzML, we used pyimzML to retrieve spectra. For the Bruker .d format, we implemented a fair random access method using SQLite coordinate mapping to eliminate sequential iteration overhead. For SpatialData/Zarr, we performed direct row slicing on the sparse CSC matrix and forced computation to dense arrays.
- **Random m/z Range Query:** To simulate ion image extraction, we generated 100 random m/z ranges (widths 100 - 1,000 bins). For imzML and Bruker formats, this required iterating through all pixels to retrieve complete spectra, reflecting the lack of columnar access. For SpatialData/Zarr, we utilized columnar slicing (matrix[:, start:end]) to aggregate intensities efficiently.
- **Random ROI Extraction:** We generated 100 random rectangular ROIs (10 - 50-pixel side lengths). To ensure a consistent computational workload across formats, we retrieved all spectra within the ROI and computed the mean spectrum. For imzML and Bruker, spectra were retrieved individually via coordinate lookups . For SpatialData/Zarr, we performed contiguous row slicing to retrieve the data matrix before computing the mean.

### Timing & Verification

All measurements used Python’s time.perf_counter(). Initialization time (file parsing and metadata extraction) was measured separately and excluded from query latencies. For lazy-loading formats (Zarr), computation was explicitly forced to prevent measurement of only query construction time. Results were aggregated from 100 iterations per pattern to report median latencies.

## Data Availability

The large-scale multi-modal spatial omics dataset generated in this study is available on Zenodo under the accession code **10.5281/zenodo.18326569**. This repository contains the corresponding data for all modalities used in **Figures 3 and 4**, including:

1. **Mass Spectrometry Imaging (MSI):** Raw Bruker .d file.
2. **Spatial Transcriptomics:** Raw Xenium output files.
3. **Histology:** High-resolution Brightfield microscopy images (.svs).

## Code Availability

Thyra is available as an open-source project under the MIT license at https://github.com/M4i-Imaging-Mass-Spectrometry/thyra. The package can be installed via PyPI.

## Acknowledgements

The authors would like to thank Dr. Veerle Melotte, Lisa Maurginther (Department of Pathology, Maastricht University) Jill Grondelaers (Maastricht University) for providing the mouse brain tissue, and assisting with the H&E, Tim Hendriks (Maastricht University) for acquiring the MALDI-MSI dataset and Ruben Jacobs (Maastricht University) for acquiring the Xenium Spatial Transcriptomics experiment.

## References

1. van Hove, E. R. A., Smith, D. F. & Heeren, R. M. A concise review of mass spectrometry imaging. Journal of chromatography A 1217, 3946–3954 (2010).

2. Korber, A., Anthony, I. G. & Heeren, R. M. Mass Spectrometry Imaging. Analytical Chemistry 97, 15517–15549 (2025).

3. Bouvier, C., Van Nuffel, S., Walter, P. & Brunelle, A. Time-of-flight secondary ion mass spectrometry imaging in cultural heritage: a focus on old paintings. Journal of Mass Spectrometry 57, e4803 (2022).

4. Vandereyken, K., Sifrim, A., Thienpont, B. & Voet, T. Methods and applications for single-cell and spatial multi-omics. Nature Reviews Genetics 24, 494–515 (2023).

5. Flinders, B. et al. Investigation into Drug-Induced Liver Damage Using Multimodal Mass Spectrometry Imaging. Journal of the American Society for Mass Spectrometry 36, 265–276 (2025).

6. Alvarez-Martin, A., Quanico, J., Scovacricchi, T., Avranovich Clerici, E., Baggerman, G. & Janssens, K. Chemical mapping of the degradation of geranium lake in paint cross sections by MALDI-MSI. Analytical chemistry 95, 18215–18223 (2023).

7. Alexandrov, T., Saez-Rodriguez, J. & Saka, S. K. Enablers and challenges of spatial omics, a melting pot of technologies. Molecular Systems Biology 19, e10571 (2023).

8. Martens, L. et al. mzML—a community standard for mass spectrometry data. Molecular & Cellular Proteomics 10 (2011).

9. Deutsch, E. Vol. 8 2776–2777 (Wiley Online Library, 2008).

10. Schramm, T. et al. imzML — A common data format for the flexible exchange and processing of mass spectrometry imaging data. Journal of Proteomics 75, 5106–5110 (2012). 10.1016/j.jprot.2012.07.026

11. Van Den Bossche, T. et al. mzPeak: Designing a Scalable, Interoperable, and Future-Ready Mass Spectrometry Data Format. Journal of Proteome Research (2025). 10.1021/acs.jproteome.5c00435

12. Abernathey, R. P. et al. Cloud-native repositories for big scientific data. Computing in Science & Engineering 23, 26–35 (2021).

13. Bhamber, R. S., Jankevics, A., Deutsch, E. W., Jones, A. R. & Dowsey, A. W. mzMLb: A Future-Proof Raw Mass Spectrometry Data Format Based on Standards-Compliant mzML and Optimized for Speed and Storage Requirements. Journal of Proteome Research 20, 172–183 (2021). 10.1021/acs.jproteome.0c00192

14. Marconato, L. et al. SpatialData: an open and universal data framework for spatial omics. Nature Methods 22, 58–62 (2025).

15. Moore, J. et al. OME-NGFF: a next-generation file format for expanding bioimaging data-access strategies. Nature Methods 18, 1496–1498 (2021). 10.1038/s41592-021-01326-w

16. metaspace-converter: Download and Convert METASPACE datasets. GitHub repository (2024).

17. Virshup, I. et al. The scverse project provides a computational ecosystem for single-cell omics data analysis. Nature biotechnology 41, 604–606 (2023).

18. Palla, G. et al. Squidpy: a scalable framework for spatial omics analysis. Nature methods 19, 171–178 (2022).

19. Hendriks, T. F. E., Eijkel, G. B., Visvikis, T., Balluff, B., Heeren, R. M. A. & Cuypers, E. One section, two worlds: single-cell integration of MALDI-MSI and spatial transcriptomics on the same single tissue section. Scientific Reports 15, 42660 (2025). 10.1038/s41598-025-26735-1

20. Heidari, E. et al. Segger: Fast and accurate cell segmentation of imaging-based spatial transcriptomics data. bioRxiv, 2025.2003.2014.643160 (2025). 10.1101/2025.03.14.643160

21. Muñoz, A. F. et al. cp_measure: API-first feature extraction for image-based profiling workflows. arXiv preprint 2507.01163 (2025).

22. Keller, M. S., Gold, I., McCallum, C., Manz, T., Kharchenko, P. V. & Gehlenborg, N. Vitessce: integrative visualization of multimodal and spatially resolved single-cell data. Nature Methods 22, 63–67 (2025). 10.1038/s41592-024-02436-x

23. Pollaris, L. et al. SPArrOW: a flexible, interactive and scalable pipeline for spatial transcriptomics analysis. bioRxiv, 2024.2007.2004.601829 (2024). 10.1101/2024.07.04.601829

